# Computational approaches for discovery of mutational signatures in cancer

**DOI:** 10.1101/154716

**Authors:** Adrian Baez-Ortega, Kevin Gori

## Abstract

The accumulation of somatic mutations in a genome is the result of the activity of one or more mutagenic processes, each of which leaves its own imprint. The study of these DNA fingerprints, termed mutational signatures, holds important potential for furthering our understanding of the causes and evolution of cancer, and can provide insights of relevance for cancer prevention and treatment. In this review, we focus our attention on the mathematical models and computational techniques that have driven recent advances in the field.

## Introduction

Cancer is a disease of the genome, in which uncontrolled clonal proliferation is initiated and fuelled by genomic alterations in somatic cells [1]. Despite the fact that a cancer genome may carry between tens and millions of somatic mutations [2,3], only a small subset of these, termed ‘driver’ mutations, are thought to be under selection and to cause neoplastic expansion [1,4]. The remaining ‘passenger’ mutations are generally believed not to confer selective advantage, and to arise from the processes involved in mutagenesis [5,6]. The collection of mutations in a somatic cell genome is the result of one or more mutational processes operating, continuously or intermittently, during the organism’s lifetime [7]. Such mutational processes include DNA damage by exogenous or endogenous agents, defective DNA replication, insertion of transposable elements, defects in DNA repair mechanisms, and enzymatic modifications of DNA, among others [8]. Many of these processes imprint a distinct pattern of mutations in the genome, known as a ‘mutational signature’ [2,9]. Therefore, the compendium of somatic changes in a cancer genome constitutes a record of the combined mutagenic effect of the specific mixture of processes moulding it [2]. Furthermore, because most mutations are passengers, they are largely beyond the effect of adaptive selection [10].

Although mutational signatures are a relatively recent concept in cancer biology, the first descriptions of genomic aberrations caused by a specific process date back to the early twentieth century, when X-rays were found to induce chromosome breakage in irradiated cells [11–13]. More-detailed mutational patterns were reported in the 1960s, notably the crosslinking of adjacent pyrimidine bases (CC, CT, TC, TT) due to ultraviolet radiation, which produces cytosine-to-thymine (C>T) and cytosine–cytosine-to-thymine–thymine (CC>TT) transitions at dipyrimidine sites [14–16]. Other causal links between mutagenic agents and patterns of somatic changes have also become established, such as the guanine-to-thymine (G>T) transversions resulting from guanine adducts that are caused by carcinogens present in tobacco smoke [17,18]. Furthermore, some chemotherapeutic agents are mutagens as well, and may imprint their own mutational signature in the cancer genomes of patients with secondary malignancies [19,20]. These examples illustrate the importance of studying somatic mutation patterns to our understanding of the molecular mechanisms of neoplasia, potentially enabling the discovery of novel mutagens [2,7,8,21]. Moreover, several authors have emphasised the potential of mutational signature analysis to provide insights of clinical significance, by informing and guiding diagnostic procedures, personalised cancer interventions and prevention efforts [19,22–27].

Recent advances in high-throughput DNA sequencing technologies have enabled studies which examine many thousands of whole cancer genomes or exomes. In parallel, new scientific avenues have been explored to identify and analyse genomic aberrations, among them the extraction of mutational signatures from collections of somatic mutations. This has produced catalogues of signatures that operate in a variety of human neoplasias [2,28–31]. While the development of methods for discovery of mutational signatures has achieved considerable success, this is still an emerging field, stemming from very recent analytical and technological breakthroughs. In this review, we aim to summarise current methodologies, in particular the mathematical models and computational techniques, which form the basis of mutational signature analysis.

## Mathematical modelling of mutational signatures

A mutational signature can be mathematically defined as a relationship between a (known or unknown) mutagenic process and a series of somatic mutation types. Many classes of genomic alterations can serve as features of a mutational signature, including singlesor di-nucleotide substitutions, small insertions and deletions (indels), copy number changes, structural rearrangements, transposable element integration events, localised hypermutation (*kataegis*), and epigenetic changes. In practice, only a limited number of features can be incorporated into the mathematical abstraction of a mutational signature, with the attention of most studies to date being focused on single-base substitutions. However, signatures based on indels [29,32] or structural variants [27,29,32] have also been described. Furthermore, certain substitution signatures are consistently associated with features such as increased numbers of indels or rearrangements of a particular class, *kataegis* events, or biases in the transcriptional strand in which mutations occur [2,28–30,33]. It is therefore useful to consider such features as biological constraints for the identification of signatures, even if precisely modelling them is more challenging.

The selected set of *K* mutation types can be expressed as a finite alphabet 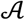, with 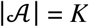, every symbol in 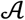 representing a distinct mutation type. This alphabet constitutes the domain of a mutational signature, which is modelled as a discrete probability density function, 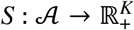. Hence, the mathematical representation of a given signature, *S_n_*, is a *K*-tuple of probability values, *S_n_* = [*s*_1*n*_, *s*_2*n*_, …, *s_Kn_*]^T^, with *s_kn_* denoting the probability of the mutation type represented by the *k*-th symbol in A being caused by the mutational process associated with *S_n_*. As probability values, the elements of *S_n_* are intrinsically nonnegative and their sum is always 1:
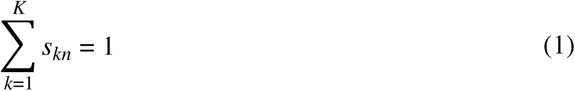

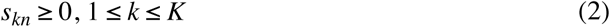

The same mutational process operating in multiple genomes may produce different numbers of mutations in each. The intensity at which a mutational process with signature *S_n_* operates in a genome *g*, expressed in terms of the number of mutations caused, is known as the ‘exposure’ to (or the ‘contribution’ or ‘activity’ of) the process, and denoted by *e_ng_*. Regarding the catalogue of somatic mutations in a cancer genome *g*, this is also defined as a vector of mutation counts over 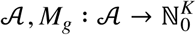, and expressed as a second nonnegative *K*-tuple: *M_g_* = [*m*_1*g*_, *m*_2*g*_,… *m_Kg_*]. (This notation of mutational catalogues, signatures and exposures will be maintained hereafter for coherence.)

A mutational catalogue can be approximately considered as a linear superposition of the signatures of the latent mutational processes that have acted at some point in the somatic cell lineage giving rise to the sampled neoplastic cells, each signature weighted by the exposure to the corresponding process. In addition, catalogues are expected to contain some level of noise arising from sequencing or analysis errors and sampling noise. Neglecting such noise, the number of mutations of the *k*-th type in the catalogue *M_g_*, *m_kg_*, can be approximated by the sum of the *k*-th element of the *N* operative mutational signatures, each weighted by its respective exposure:
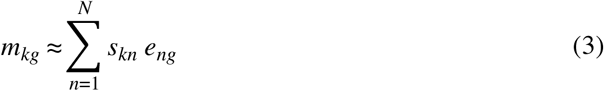

Most of the existing mathematical approaches to mutational signature inference have focused on single-base substitutions as mutation features, maintaining the convention established by Nik-Zainal *et al.* [33] and Alexandrov *et al.* [2]. In this scheme, substitutions are first classified into six categories, by representing the change at the pyrimidine partner in the mutated base pair (e.g. a guanine-to-adenine substitution, G>A, is instead expressed as a cytosine-to-thymine change, C>T, in the complementary strand). This classification is then extended by considering the immediate sequence context of the substitution, usually the adjacent 5’ and 3’ bases. The six substitution types are thus translated into 96 trinucleotide mutation types (6 substitution types × 4 types of 5’ base × 4 types of 3’ base). An extensive literature supports the need for at least a trinucleotide context of mutations in order to distinguish the mutational patterns induced by a variety of mutagens. In addition, there have been attempts to deconvolute signatures using a five- or seven-base sequence context, resulting in 1536 and 24,576 mutation types, respectively [27,34,35]. Further elaboration can also be achieved by considering the transcriptional strand of mutations in transcribed regions. Nevertheless, expanding the range of mutation types normally implies a decrease in the observed number of mutations per type, which may curb the power to identify patterns.

In a generalisation that considers *N* different mutational processes acting in a collection of *G* cancer genomes, with mutational catalogues defined over *K* mutation types, the catalogues, signatures and exposures can be mathematically expressed as matrices named *M*, *S* and *E*, respectively (**Fig. 1a**):

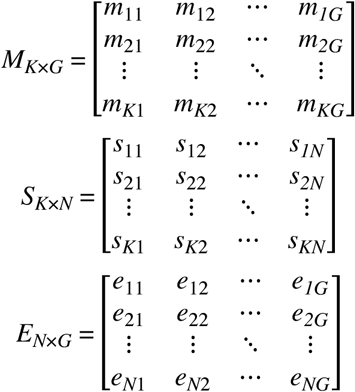

**Fig. 1.**
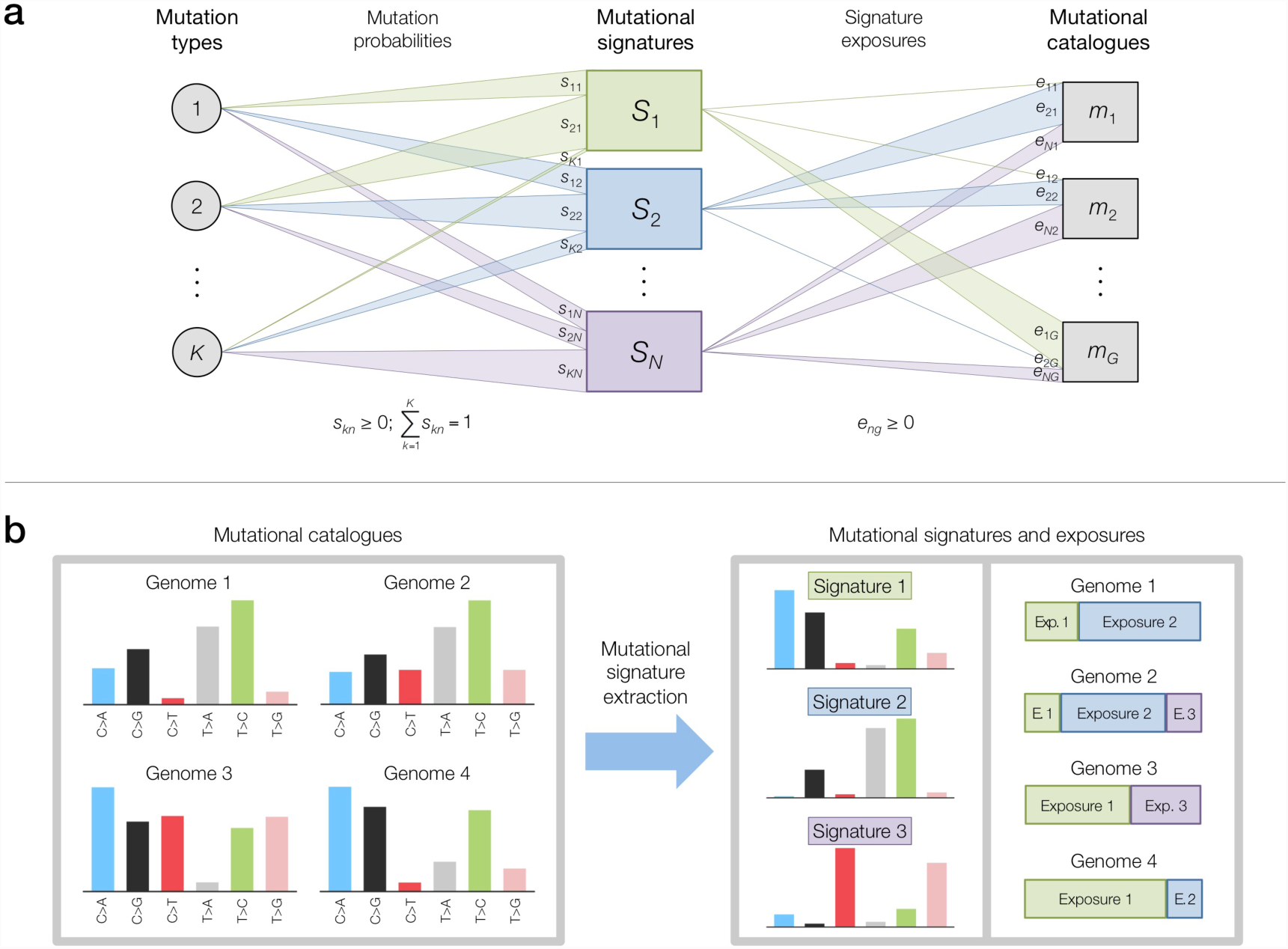
Mathematical modelling and deconvolution of mutational signatures. **(a)** Diagram illustrating the modelling of mutational signatures as probabilistic relationships between mutation types and mutational processes operative in genomes, for a general case with *K* mutation types, *N* mutational processes and *G* genomes. The notation of signatures, exposures and mutational catalogues follows that used in the main text. The varying widths of the links between mutation types and signatures (mutation probabilities), and between signatures and catalogues (signature exposures) represent the observation that varying values of *s_kn_*and *e_ng_* reflect the specific mutational profile of each signature and the exposure composition of each genome. Nonnegativity constraints for mutation probabilities and signature exposures are specified directly below them. **(b)** Example of *de novo* signature extraction, for a case with *K* = 6 mutation (single-base substitution) types, *N* = 3 mutational signatures and *G* = 4 mutational catalogues. Starting from the set of catalogues (depicted here as mutational profiles, each bar corresponding to a distinct mutation type), *de novo* extraction methods determine the set of mutational signatures (represented as consensus mutational profiles) and exposures (depicted here as proportions of the mutations in each catalogue, for simplicity) that reconstruct the original mutational catalogues with minimal error.

Consequently, the approximate description of a mutational catalogue as a sum of signatures multiplied by their exposures, expressed in (3), is generalised into matrix form:
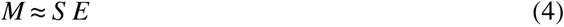

By adopting this mathematical representation, the problem of inferring the mutational signatures and exposures that best account for a given collection of observed catalogues becomes equivalent to finding the instances of *S* and *E* that reproduce *M* with minimal error. This is, in turn, connected to the problem of determining the number of signatures, *N*, that optimally explains the data in *M* (**Fig. 1b**). This process is sometimes referred to as *de novo* extraction, inference, deciphering, or deconvolution of mutational signatures. By contrast, the simpler problem of signature refitting is characterised by both *M* and *S* being known *a priori*.

## Computational approaches for mutational signature discovery

A host of computational strategies have been advanced to tackle the problem of signature discovery as formulated above; these are presented below and summarised in **Table 1**.

**Table 1.** . Published software packages for mathematical inference of mutational signatures. (Abbreviations: EM: expectation–maximisation; MCMC: Markov chain Monte Carlo; NMF: nonnegative matrix factorisation; WTSI: Wellcome Trust Sanger Institute.)

| Software | Mathematical framework | *De novo* signature extraction | Incorporation of mutational opportunity | Notable aspects | Programming language(s) | Reference(s) |
| --- | --- | --- | --- | --- | --- | --- |
| WTSI Mutational Signature Framework | NMF | Yes | No | • First mathematical model of signatures<br>• Extensive development and application<br>• ‘Gold standard’ status | MATLAB | [34] |
| SomaticSignatures | NMF | Yes | No | • Ease of use<br>• Integration in Bioconductor | R | [48] |
| MutSpec | NMF | Yes | No | • Ease of use<br>• Graphic user interface | R, Perl (Galaxy platform) | [57] |
| EMu | Probabilistic (EM, Poisson model) | Yes | Yes (tumour-specific) | • First probabilistic model of signatures<br>• First modelling of mutational opportunity<br>• Automatic estimation of number of signatures | C++ | [61] |
| BayesNMF | Bayesian NMF (Poisson model) | Yes | No | • Automatic estimation of number of signatures | R | [70,71] |
| signeR | Bayesian NMF (MCMC EM, Poisson model) | Yes | Yes (tumour-specific) | • Automatic estimation of number of signatures<br>• Differential exposure analysis<br>• Unsupervised sample classification | R, C++ | [74] |
| pmsignature | Probabilistic (EM, independent model) | Yes | Yes | • Simplified mathematical model<br>• Increased number of signature features<br>• Alternative visual representation | R, C++ | [35] |
| deconstructSigs | Multiple linear regression | No | Yes | • Analysis of signature activities in individual tumours | R | [86] |

### Nonnegative matrix factorisation

The unsupervised learning technique of nonnegative matrix factorisation (NMF) [36,37] was devised to explain a set of observed data utilising a set of components, the combination of which approximates the original data with maximal fidelity. NMF is distinguished from similar techniques, such as principal component analysis (PCA) or independent component analysis (ICA), in that nonnegativity is enforced for the values composing both the components and the mixture coefficients, and that no orthogonality or independence constraints are imposed (therefore permitting partially or entirely correlated components). These features make NMF especially well-suited to the problem of mutational signature inference, because of the intrinsic nonnegativity of the matrices in the mathematical model presented above. Moreover, NMF has repeatedly stood out as a powerful technique for the extraction of meaningful components from various types of high-dimensional biological data [38–42], besides successful applications in other fields [39].

NMF constituted the basis of the first computational method for mutational signature inference, the **Wellcome Trust Sanger Institute (WTSI) Mutational Signature Framework** (hereafter referred to as the **WTSI Framework**). This was published, together with the mathematical model introduced above, in a landmark work by Alexandrov *et al*. [34], which enabled the first detailed delineations of mutational signatures in human cancer [2,33,43]. The WTSI Framework performs NMF on a set of mutational catalogues by building upon an implementation, developed by Brunet *et al*. [38], of the multiplicative update algorithm devised by Lee and Seung [36,44]. More formally, given a set of mutational catalogues, *M*, composed of *G* genomes defined over *K* mutation types, the method extracts exactly *N* mutational signatures (with 1 ≤ *N* ≤ min{*K*, *G*} – 1), by finding the matrices *S* and *E* that approximately solve the nonconvex optimisation problem derived from (4), with the selected matrix norm being the Frobenius reconstruction error:
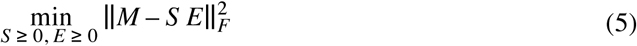

The algorithm first initialises *S* and *E* as random nonnegative matrices, and reduces the dimension of *M* by removing those mutation types that together account for ≤1 % of all the mutations. Two steps are then iteratively followed: (*a*) Monte Carlo bootstrap resampling of the reduced catalogue matrix, and (*b*) application of the multiplicative update algorithm to the resampled matrix, finding the instances of *S* and *E* that minimise the Frobenius norm in (5). After completion of the iterative stage, partition clustering is applied to the resulting set of signatures, in order to structure the data into *N* clusters. The *N* consensus signature vectors, which compose the averaged signature matrix, 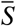, are obtained by averaging the signatures in each cluster. Since each signature is related to a specific exposure, the averaged exposure matrix, 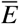, can be inferred from 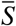. In cases where the mutational catalogues have been derived from cancer exomes, the extracted mutational signatures should thereafter be normalised to the trinucleotide frequencies of the whole genome.

The WTSI Framework requires the number of signatures to infer, *N*, to be defined as a parameter. Because the number of signatures present in the data is normally not known *a priori*, the framework needs to be applied for values of *N* ranging between 1 (or the smallest plausible number of signatures) and min{*K*, *G*} – 1. For each value of *N*, the overall reproducibility (measured as the average silhouette width [45] of the signature clusters, using cosine similarity) and Frobenius reconstruction error are calculated, and the best value is selected such that the resulting signatures are highly reproducible and exhibit low overall reconstruction error. Nevertheless, the manual determination of *N* on the basis of these criteria is perhaps the most heavily criticised aspect of the WTSI Framework. Accurate estimation of the number of mutational signatures, besides remaining one of the thorniest facets of mutational signature analysis, is crucial given the associated risks of inferring signatures that merely describe the noise in the data by overfitting (through overestimation of *N*), or insufficiently separating signatures present in the data by underfitting (through underestimation of *N*).

Although the NMF approach has proven highly effective, especially when applied to large cohorts of cancer genomes, it is not without conceptual limitations [34]. The first of these lies in the number of catalogues required, which is a limiting factor on the number of signatures that can be accurately extracted, and rises exponentially with *N*. The number of mutations per catalogue also influences the power to infer signatures, with a small set of densely mutated genomes being more informative than a large number of sparsely mutated genomes. In fact, the influence of catalogues with extreme mutation burdens (hypermutated genomes) on the NMF process can hinder the detection of signals from less-mutated catalogues. Furthermore, mutational signatures exhibiting higher exposures can generally be identified more easily and accurately. Sensitivity to initial conditions is another major limitation, arising from the high dimensionality and inherent nonconvexity (presence of multiple local minima) of the optimisation problem posed by (5). This aspect of NMF has attracted particular attention in the past, leading to the proposal of alternative initialisation strategies [46,47] that might outperform the random initialisation adopted by the WTSI Framework.

In more recent analyses, the WTSI working group has significantly refined their own application of the WTSI Framework, in order to enhance power and accuracy; however, such refinements have not been incorporated in the publically available software. Firstly, an additional analysis step can follow the deconvolution of consensus mutational signatures, which centres on precisely estimating the contribution of each signature to each genome [28]. This is individually achieved for each catalogue through minimisation of a variation of the function shown in (5); the difference lies in *S* now being known, and harbouring only the consensus mutational patterns of the processes that operate in the tumour type of the sample (these are known from the signature extraction process). Notably, additional biological constraints are imposed in the selection of the processes included in *S*; these require that, for each candidate process, at least one associated genomic feature (e.g. transcriptional strand bias or enrichment in aberrations of a specific type) be present in the examined sample. The second enhancement consists of a ‘hierarchical signature extraction’ process [29], which is directed to increase the power to identify signatures exhibiting either low activity or limited representation across the sample cohort. Here, the WTSI Framework is initially applied to the original matrix, *M*, containing all the somatic catalogues. After identification of signatures, those samples that are well-explained by the resulting mutational patterns are removed from *M*, and the method is re-applied to the remaining catalogues. The process is repeated until no new signatures are discovered, and the additional step for estimating signature contributions described above is then applied to all the consensus patterns.

Following the success of the WTSI Framework, other software tools have been released that exploit NMF to decipher mutational signatures. The **SomaticSignatures** package, developed by Gehring *et al*. [48], provides an R implementation of the NMF algorithm by Brunet *et al*. [38]. It aims to offer a more accessible approach to signature inference, featuring additional normalisation and plotting routines and allowing integration with widely used Bioconductor [49] workflows and data structures. On the other hand, this accessibility is accompanied by a notable shortage of options for fine-tuning of the inference process. In addition, the package allows the application of PCA for *de novo* signature extraction; however, since it does not enforce nonnegativity, PCA is implausible from a biological standpoint, and unlikely to be fruitful. Despite this, and due to its simplicity and adherence to the Bioconductor framework, SomaticSignatures has become the tool of choice in a number of recent cancer studies [50–56].

**MutSpec** is a third framework, presented by Ardin *et al*. [57], that exploits NMF through the R package developed by Gaujoux and Seoighe [58]; this provides an interface to several NMF implementations, including that by Brunet *et al*. [38]. Moreover, MutSpec stands out for being the first published tool in the field that features a comprehensive graphical user interface, with a view toward empowering a wider variety of researchers, including those with limited bioinformatics expertise, to perform analyses of mutational catalogues. MutSpec accomplishes this by building upon the open-source Galaxy platform [59,60], which allows integration of multiple bioinformatics tools in an accessible and reproducible manner.

Although both SomaticSignatures and MutSpec ultimately apply the same implementation of the multiplicative update algorithm for NMF [38] originally adopted by the WTSI Framework, it should be noted that these packages may not produce identical results to those of the latter, since they lack the computationally intensive pre-processing and bootstrapping routines that complement the application of NMF in the method devised by Alexandrov *et al*. [34]. Nevertheless, SomaticSignatures and MutSpec do adopt the definition of mutational signatures as probability vectors over single-base substitution types in a trinucleotide context. It is worth noting that one recent study [27] that applied both the WTSI Framework and SomaticSignatures for *de novo* extraction of signatures from esophageal adenocarcinoma genomes reported a high similarity between the core mutational patterns identified by both tools.

### Expectation–maximisation

In contrast to the numerical optimisation approach to mutational signature inference expressed by (5), probabilistic frameworks have also been devised which exploit the intrinsically stochastic nature of mutagenesis. These frameworks have been claimed to be better-suited to deal with mutational stochasticity, which is partly responsible for the noise observed in mutational catalogues and becomes more prominent as less-mutated genomes, or smaller genomic regions, are examined.

The first probabilistic approach in the field was developed by Fischer *et al*. [61], under the name **EMu**. It builds upon the insight that the NMF optimisation problem posed by the WTSI Framework can be recast as a probabilistic model, in which the observed mutation counts (*M*) are distributed as independent Poisson random variables, parameterised by the product of the matrices of signatures (*S*) and exposures (*E*). Given some assumptions, such as that the quantity being minimised in NMF is a type of Bregman divergence [62], the two approaches are equivalent [63–65]. Estimation of *S* and *E* is performed through an expectation–maximisation (EM) algorithm [66]. Notably, the probabilistic setting also addresses the determination of the most plausible number of signatures, *N*, as a model selection problem.

Another novelty of EMu is the incorporation of tumour-specific variation in mutational opportunity across different sequence contexts. Mutational opportunities, which derive from the sequence composition of a genome, can be expressed as a nonnegative *K-*tuple containing the opportunity for each mutation type in the genome *g*, *O_g_* = [*o*_1*g*_, *o*_2*g*_, …, *o_Kg_*]. For single-base substitutions in a trinucleotide context, the opportunities correspond to the frequencies of each trinucleotide type in each genome. Explicitly accounting for the opportunity for mutations to occur is especially relevant given that the relative frequency of certain sequences in the human genome (e.g. underrepresentation of CpG dinucleotides) can exert undesired biases on the inferred mutational patterns. In addition, copy number alterations, which are frequent in cancer genomes [1,67], can substantially alter the mutational opportunity in affected regions across tumours. The divergence in sequence composition across genomic segments also makes opportunity a relevant factor in the determination of signature contributions in a specific region. The probabilistic framework and explicit dependence on opportunity are intended to increase adaptability for the analysis of signatures in short genomic regions.

Fischer *et al.* make use of a Poisson-distributed probabilistic model to describe the mutational catalogue of a given genome as the result of a stochastic process of mutation accumulation. Assuming the *N* mutational processes to be mutually independent, the probability of observing the catalogue *M_g_* = [*m*_1*g*_, *m*_2*g*_, …, *m_Kg_*] is given by:
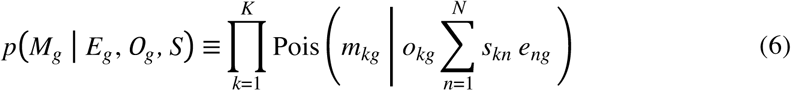

In this model, the mutational signatures, *S*, act as the shared model parameters, and the signature exposures, *E*, as the hidden data. The end of the EM procedure is to find maximum likelihood estimates of both, thereby solving the deconvolution problem. The algorithm starts by making an initial guess of the model parameters, *S*^(0)^, and thereafter iterates through two steps. In the first, denoted E-step, an estimate is obtained for the signature exposures, 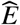, given the current parameter guess, *S*^(*k*)^. In the subsequent M-step, 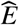 is used to update the parameter estimate for the next iteration, *S*^(*k*+1)^. Iteration through these steps finishes when the likelihood of the observed data, *p*(*M|S*), converges to a local maximum.

The data likelihoods obtained for different values of *N* are compared in order to determine the number of mutational processes involved. Because increasing *N* normally leads to a better explanation of the data, due to the higher number of available model parameters, the likelihood generally rises with *N*. Overfitting of the data is avoided applying the Bayesian information criterion (BIC) [68], a model selection criterion whose second term corrects for the model complexity:
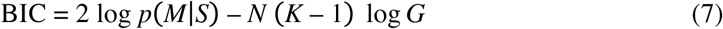

The BIC is calculated for each of the models, and the one exhibiting the highest BIC value is selected [68,69]. After inference of signatures, EMu can estimate both the global exposures in each genome and the local exposures per genomic region. Inference of local exposures is performed by dividing each genome into non-overlapping segments of equal length, and using the estimated global exposures as an informed prior distribution. The patterns of variation in local exposures can subsequently be compared within and across genomes.

It is worth noting that, while EMu builds upon a valid alternative interpretation of NMF, which considers the latter as an application of EM to a particular problem [64], the novel concepts and advantages of the method presented by Fischer *et al.* are not intrinsic properties of the EM paradigm, but explicit enhancements that are amenable to assimilation by other approaches. On the other hand, EMu suffers from the same sensitivity to initial conditions as conventional NMF, and it may as well benefit from alternative initialisation strategies. Despite this, EMu successfully exploits a probabilistic formulation of mutational signature inference to address previously unexplored aspects, namely the incorporation of context- and tumour-specific opportunity for mutations, the estimation of local signature exposures, and the direct determination of the number of mutational processes.

### Bayesian NMF

As noted above, the WTSI Framework has been criticised for requiring a manual selection of the number of mutational signatures, *N*, on the basis of heuristics that are indicative of the goodness of the solutions. While EMu addresses this issue by means of a purely probabilistic methodology, alternative approaches have proceeded by wrapping NMF in a Bayesian framework, partly with a view toward improving estimation of *N*.

The **BayesNMF** software by Kasar *et al.* [70] and Kim *et al.* [71] is based upon a variant of NMF proposed by Tan and Févotte [72]. Similarly to the strategy introduced by Fischer *et al.* [61], BayesNMF exploits the compatibilities between NMF and a Poisson generative model of mutations. More specifically, the number of mutations of the *k*-th type in a genome *g*, *m_kg_*, is assumed to be the combination of *N* independent mutation burdens, 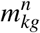 (with 1 ≤ *n* ≤ *N*); such burdens are in turn assumed to be generated by a Poisson process parameterised by mutation-type- and genome-specific rates, such that the expected number of mutations attributed to signature *S_n_* is:
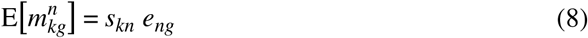

The properties of the Poisson process [73] then imply that *m_kg_* is also Poisson-distributed as:
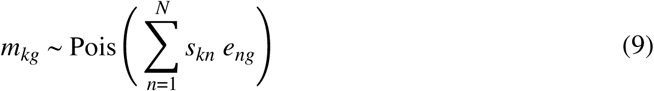

Consequently, as already seen, the estimation of signatures (*S*) and exposures (*E*) by maximising the likelihood of the observed data (*M*), given the expectation E[*M*] = *S E*, is equivalent to the minimisation of a particular Bregman divergence [62] between *M* and the matrix product *S E* through NMF [72]. However, BayesNMF addresses the selection of *N* implicitly through a technique known as ‘automatic relevance determination’ [72], which ‘prunes’ or ‘shrinks’ those components in *S* and *E* which are inconsequential, not contributing to explaining *M*. Each signature *S_n_* is therefore assigned a relevance weight, *W_n_*; then, after imposing appropriate priors on the parameters, NMF inference is performed via numerical optimisation. During this process, the columns of *S* and rows of *E* corresponding to inconsequential pairs of signatures and exposures are shrunk to zero by their relevance weights. The effective dimensionality, corresponding to the estimated number of mutational signatures, is given by the final number of nonzero components.

Notably, the authors have extended their method to explicitly incorporate the transcriptional strand of mutations [71], resulting in a model with 192 trinucleotide mutation types (96 for each strand). While the WTSI Framework does not explicitly account for transcriptional strand biases, some studies have used this and other genomic features as biological constraints for validating the presence of specific signatures in a sample [28]. Moreover, models incorporating transcriptional strand information are only suitable for mutations in transcribed regions.

Another notable aspect of the application of BayesNMF, particularly that presented by Kim *et al.* [71], is the manner in which the excessive influence of hypermutated catalogues on the inference is moderated. This is based on equally partitioning the mutations in hypermutated genomes into multiple artificial catalogues, which maintain the mutational profile of the original tumour. The number of artificial catalogues is chosen such that their contribution becomes similar to that of non-hypermutated samples, without altering the overall number of mutations. Because of the linear properties of NMF [36], the number of mutations attributed to each signature in the original genomes can be reconstructed by summing the exposures in their respective artificial catalogues. As a measure to overcome sensitivity to initial conditions, Kim *et al.* [71] also performed multiple applications of the method with random initial conditions.

A second Bayesian approach to NMF has been recently proposed by Rosales *et al.* [74] in the form of the **signeR** package. This follows an empirical Bayesian approach to NMF which considerably differs from the strategy devised by Kasar *et al.* [70] and Kim *et al.* [71]. Firstly, the authors account for tumour-specific mutational opportunities, following the example set by Fischer *et al.* [61]. The number of mutations of the *k*-th type in a genome *g*, *mk_g_*, is assumed to be a Poisson-distributed variable, with a rate incorporating the mutational opportunity, *o_kg_*:
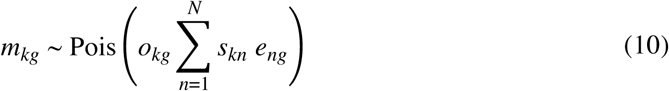

The matrices *S* and *E*, which are the parameters of the generative Poisson process, are initialised either by sampling from their (Gamma) prior distributions, or by applying numerical NMF via the implementation developed by Gaujoux and Seoighe [58]. The central method for inference is based on a combination of Markov chain Monte Carlo (MCMC) and EM techniques, which are applied in an iterative fashion [75]. This MCMC EM strategy provides a posterior distribution of the NMF model, from which estimates for the mutational signatures and exposures can be derived. The MCMC EM algorithm, in which the chosen MCMC variant is a Metropolised Gibbs sampler, is applied to obtain a series of MCMC samples from the posterior distributions of the model parameters (*S* and *E*), hyperparameters and hyperprior parameters. These samples can be subsequently used to derive point estimates and posterior statistics for signatures and exposures. Estimation of the number of mutational signatures is tackled, as in EMu, by means of the BIC, which is described in (7) and computed as the median of the BIC values across the MCMC samples.

In addition to this Bayesian NMF framework, Rosales *et al.* [74] introduce two novel applications of the method. The first is the incorporation of an *a priori* categorisation of samples, on the basis of independent knowledge (e.g. clinical data), in order to determine whether the exposure of any of the mutational signatures diverges significantly between the defined categories. Secondly, a measure known as ‘differential exposure score’, which results from this analysis of exposures, can be used to assign unclassified samples to one of the categories, using a *k*-nearest neighbours algorithm [76]. This ability for unsupervised clustering of tumours may prove especially relevant for clinical cancer prognosis.

### Independent probabilistic model

An unconventional approach to mutational signature discovery, which stands out for the adoption of a novel probabilistic model of signatures, has been introduced in the **pmsignature** R package by Shiraishi *et al.* [35]. Their model is termed ‘independent’ because, in contrast to the conventional ‘full’ model employed by all other methods, it decomposes mutational signatures into separate features (such as substitution type, flanking bases or transcriptional strand bias), which are assumed to be mutually independent. The notion of independence across features of a signature, if counterintuitive, simplifies the model drastically by reducing the number of parameters per signature. This, in turn, allows incorporation of additional signature features, such as extended sequence context. For instance, the mutational pattern defined by single-base substitutions in a pentanucleotide sequence context results in *K* = 1536 mutation types, or 1535 free parameters per signature, in the full model. Generally, accounting for the *n* adjacent bases 5’ and 3’ of the mutated site results in (*K* – 1) = (6 × 4^2*n*^ – 1) free parameters in the full model. This imposes a practical limit on the number of features that can be incorporated into a signature, because both inference stability and interpretability of the inferred signatures decline as the parameter space gains in dimensionality. The consequence is a constrained flexibility of full models; these, for example, normally consider only a trinucleotide sequence context, thus ignoring the information potentially harboured by farther adjacent nucleotides [77,78].

The work of Shiraishi *et al.* [35] can be seen as a quantum leap in the modelling of mutational signatures. Instead of belonging to a single mutation type, each mutation is modelled as having *L* distinct features, each with its own range of discrete values, and is therefore represented by a feature vector of length *L*. A signature *S_n_* is characterised using an *L*-tuple of parameter vectors, *F_n_* = [*f_n_*_1_, *f_n_*_2_,…,*f_nL_*], where *f_nl_* is the probability vector of the *l*-th feature in signature *S_n_*, its length being equal to the number of possible values of the feature. In this model, single-base substitutions on a pentanucleotide context are represented using five features (substitution and four flanking bases). Each feature being an independent probability vector, this involves (6 – 1) + 4 × (4 – 1) = 17 free parameters, instead of 1535. In general, incorporating the *n* adjacent bases on each side of the mutated site requires only (5 + 6*n*) parameters. Remarkably, this independent model of signatures can be considered as a generalisation of the full model; the latter would be the simplest case of independent model, where all the signature features have been collapsed into a single attribute, the ‘mutation type’, which contains all the possible feature combinations.

Instead of using numbers of mutations, pmsignature models the contribution of a signature as the proportion of mutations attributed to it in each genome. Such proportions, denoted by *q_gn_*, are termed ‘membership parameters’, due to the close relationship between this model of mutations and the so-called mixed-membership or admixture models [79] (also known as latent Dirichlet allocation models [80]), which have been extensively applied to population genetics and document clustering problems. In pmsignature, each mutation is assumed to be the result of a two-step generative model: first, a mutational signature is selected according to the membership parameters of the current catalogue; second, the features of the mutation are generated according to the multinomial distribution described by the chosen signature. Of note, informative parallelisms between NMF and admixture models have been previously noted by other authors [81], suggesting that current methods could benefit from the experience gained in applications of the latter.

The central parameters of the independent model, namely the sample membership proportions, *q_gn_*, and the signature parameters, *F_n_*, need to be estimated from the observed catalogues; this is done by means of an EM algorithm [66]. In order to account for the tendency of EM to converge to different local maxima depending on the initial conditions, the algorithm is applied on multiple initial configurations, before choosing the solution that exhibits maximum likelihood overall. To model mutational opportunity, instead of using probabilistic coefficients, pmsignature employs a ‘background signature’ corresponding to the genome frequencies of the types of nucleotide association considered (e.g. pentanucleotides). However, this background signature is based on the human reference genome, thus negating incorporation of sample-specific variegation in opportunity. Regarding the estimation of the number of mutational processes, an analogous strategy to that implemented by Alexandrov *et al.* [34] is adopted, with *N* being manually chosen such that the likelihood is sufficiently high, and the standard errors of the parameters are sufficiently low. In addition, *N* is selected such that the resulting set of mutational signatures does not contain any pair of signatures which seem to correspond to the same mutational process (signatures exhibiting similar feature patterns and membership parameters). Hence, a more versatile strategy to automatically determine *N* would constitute a major improvement of the method.

The consequence of adopting a simpler model in pmsignature, as reported by the authors [35], is a gain in power and stability, which allows inference of more-accurate and - reproducible signatures from smaller sample cohorts. Moreover, the reduction in parametric complexity enables the incorporation of additional contextual features, such as extended sequence context, transcriptional strand, copy number and epigenetic states. The consequent gain in signature resolution can potentially prompt the unveiling of novel mutational patterns and associated biological insights. Nevertheless, it must be noted that the estimation of signature parameters in pmsignature is severely impaired by its disregard of those mutational patterns lacking distinctive features (known as uniform or ‘flat’ signatures). The set of mutational signatures extracted by Shiraishi *et al*. [35] from previously published data does not feature any such ‘flat’ signature, even when some of these have been identified in the same data set [2] and experimentally validated by other groups [82,83]. The inability to detect signatures with uniform patterns therefore undermines the potential of this model in its current form.

In order to simplify the visualisation of signatures with multiple features, the authors have also introduced a novel graphical representation [35], closely related to sequence logos [84], that provides a schematic view of the distinctive characteristics of a signature. Albeit reliant on the illustration of probabilities as surface areas, which are often difficult to interpret visually [85], diagrammatic representations of this kind will likely become indispensable if the resolution of signatures is to be significantly enhanced, since the interpretation of mutational patterns expressed as plain probability distributions would soon become impractical.

### Mutational signature refitting

From the perspective of the NMF model, the problem of refitting mutational signatures consists of estimating the exposures (*E*) of a given set of signatures (*S*) in a collection of mutational catalogues (*M*), with the actual number of operative processes (*N*) being known or unknown. Because *S* is known *a priori*, signature refitting is a much more tractable problem than *de novo* signature inference. In consequence, signature refitting does not suffer the requirement of large sample cohorts to achieve power and accuracy, being even applicable to individual genomes.

The **deconstructSigs** R package, recently developed by Rosenthal *et al.* [86], is currently the only published method explicitly designed for mutational signature refitting. It adopts an iterative multiple linear regression strategy to estimate the linear combination of signatures that optimally reconstructs the mutational profile of each genome in *M*, imposing nonnegativity on the inferred signature exposures. Mutational catalogues are modelled as mutation proportions, instead of counts, and normalisation by mutational opportunity is enabled through the incorporation of the trinucleotide frequencies from the reference human genome. The iterative fitting algorithm, which is applied separately to each catalogue, starts by discarding those signatures in which a mutation type that is absent from the examined catalogue has a probability above 0.2. This prevents consideration of signatures that, according to their mutational profiles, are unlikely to be present in the tumour. An initial signature is then selected, such that the sum of squared errors (SSE) between the signature and the mutational profile of the catalogue is minimised. The exposure value that minimises the SSE for the chosen signature is set as the only positive exposure. In successive iterations, each of the remaining signatures is evaluated to find the exposure value that minimises the SSE between the reconstructed profile (including the previously incorporated exposures and the candidate one) and the mutational profile of the tumour. The signature achieving minimum SSE is selected, and its optimal exposure is incorporated to the reconstructed profile. The process continues until the difference in SSE before and after an iteration falls below an empirically determined threshold of 10^−4^; the estimated exposures are then transformed to proportions. Finally, any exposure lower than 0.06 (6%) is discarded, in order to exclude spurious signatures; this minimum exposure threshold was also empirically determined from simulation studies.

An iterative regression strategy has important associated risks, the most prominent being the impossibility of reducing or removing the contribution of a signature after it has been selected. Consequently, a signature that is actually absent from the sample might be unalterably chosen in the initial iterations, only because it fits the overall profile of the tumour better than any other signature. This is not a rare situation, since one-third of the currently published mutational signatures [31] (all of which are by default included in *S*) are mostly composed of cytosine-to-thymine (C>T) changes. Thus, for example, a mutational profile arising from the combination of two given signatures may initially be best fitted by a third signature which does not actually contribute to the mutational profile, but which significantly resembles it. Two measures to minimise the risk of misfitting are: (*a*) carefully selecting the signatures to include in *S*, preferring those that have been already associated with the examined tumour type; and (*b*) considering knowledge about additional genomic features linked to the activity of a mutational signature in a genome. Limiting the set of candidate signatures also lessens the risk of overfitting, especially given that the number of signatures, *N*, is indirectly determined in this method through the empirically set thresholds for change in SSE and minimum exposure value. On the other hand, such measures increase the opportunity for the biases of the investigator to influence the outcome.

Despite such concerns, the identification of mutational signatures in individual tumours through refitting harbours extreme potential, as emphasised by Rosenthal *et al.* [86] and demonstrated by the number of studies that have adopted their method in the short time since its publication [54,87–90]. When used for refitting well-validated signatures in specific cancer types, deconstructSigs has the power to detect mutational processes that operate only in small subsets of genomes, without the complexity or requirement of large cohorts that characterise *de novo* approaches. Some remarkable applications are the comparison between processes operative across different cancer subtypes, and the analysis of variegation in signature activities over time within a single tumour, or between primary and metastatic sites in a same patient. As genomic examination of individual malignancies is gradually incorporated into clinical practice, a straightforward method to ascertain which mutational processes operate in a cancer genome, and to what extent, potentially including their temporal and spatial evolution, will constitute an invaluable instrument for the advancement of personalised cancer therapy.

### Alternative approaches

Apart from the ones described here, both *de novo* inference and refitting of mutational signatures are amenable to many other computational approaches, including purely Bayesian techniques (e.g. hierarchical Dirichlet processes), global optimisation metaheuristics (e.g. simulated annealing), and nonlinear optimisation algorithms (e.g. sequential quadratic programming). When considering the design of novel methods for the analysis of mutational signatures, the special properties of each technique, such as propensity for overfitting, sensitivity to initial conditions, computational cost and scalability, should be thoughtfully considered. In the near term, fresh methodologies are likely to arise which build upon either the mathematical models of signatures already developed, or entirely new ones. Furthermore, because signature refitting poses a much simpler mathematical problem than *de novo* signature deconvolution, approaches based on well-established mathematical or statistical paradigms could be implemented with little effort, as substantiated by works that have already accomplished signature refitting through some of the aforementioned techniques [27,91,92].

## Discussion

In the relatively short time since its first reported application [33,43], the deconvolution of mutational signatures has proven a successful analytical technique. Numerous authors have highlighted the potential of mutational signature analysis in the settings of cancer treatment and prevention. The proposed applications thus far include the use of signatures (*a*) as genetic biomarkers of early malignancy or exposure to carcinogenic agents, especially in combination with ‘liquid biopsy’ diagnostic techniques [23,26]; (*b*) to stratify patient cohorts into subgroups indicative of distinct dominant aetiological factors, with the aim of suggesting targeted therapies that may benefit some subgroups, on the basis of the molecular mechanisms involved [19,22,24,27,93]; (*c*) to discover or support causative links between exposure to known or novel carcinogens and the development of particular cancer types, by determining the extent to which those carcinogens contribute to mutagenesis [25,26,94,95]; (*d*) to evaluate the safety of chemotherapeutic agents, some of which have been shown to contribute to the mutation burdens in exposed patients [19,20]; (*e*) to drive novel molecular research directed at establishing links between mutagens or molecular processes and currently unexplained (‘orphan’) signatures [19], or to tease apart the individual fingerprints hidden in composite mutational patterns, such as that of the complex chemical mixture in tobacco smoke [26]; and (*f*) to contribute toward public awareness and education of the cancer risk associated with preventable exposures to certain mutagens (currently, mainly tobacco smoke, ultraviolet light, aristolochic acid, aflatoxin B1 and some pathogen infections) [2,25,26,94,95].

From a biological standpoint, the potential of mutational signature analysis to identify and quantify the contributions of mutagenic processes operative in cancer genomes makes it an outstanding tool for further delving into the fundamental causes and mechanisms of tumorigenesis [7,95]. For instance, by contrasting the mutational mechanisms that operate in normal and cancer genomes, the study of signatures has helped to settle the long-standing debate around whether the mutation rates and processes shaping the genomes of normal cells can account for the aberrations found in cancer genomes [23,96].

The WTSI Mutational Signature Framework, with a considerable number of successful applications in large-scale genomic studies of cancer [2,22,24,25,27–30,32,33,43,94,97], represents the current state-of-the-art of the NMF approach to signature deconvolution. Consequently, it acts as a *de facto* ‘gold standard’ in the field. In spite of this, the method has several conceptual limitations, especially the requirement of extensive cohorts of genomes, and harbours potential for further methodological refinements [34]. Different enhanced flavours of NMF have been proposed [46,72,98–106] which might hold the key to improving the effectiveness of the WTSI Framework’s model, for example by incorporating additional sparsity constraints. Other distinct statistical approaches to signature inference have been proposed with a view towards overcoming the limitations of conventional NMF, which turn to either Bayesian approximations to NMF [71,74] or entirely probabilistic models [35,61,86]. Interestingly, independent works [25,27] have performed direct comparisons between some of these methods and reported notable coherence between their outcomes, in spite of their divergent mathematical frameworks. Other approaches, while still adhering to the classic NMF formulation, intend to facilitate signature analysis by means of user-friendly graphical interfaces [57] or integration in popular bioinformatic frameworks [48]. As a mounting number of medium-scale studies aspire to probe the mutational mechanisms operating in specific cancer types or subtypes, methods that enable simple and accurate analysis of signatures are definitely welcome contributions to the field.

The identification of mutational signatures in cancer genomes remains a daunting endeavour, despite the breakthroughs it has spurred. In the short term, some of the computational strategies reported here will likely be subjected to significant refinement, or extended through the release of new software, while fresh approaches to signature discovery, using yet-unexploited techniques, are also sure to arrive. In the longer term, it must be noted that current methods base their signature models exclusively on mutational profiles, and fail to incorporate other experimental and clinical knowledge about mutational processes. Instead, current studies rely on a manual, informal consideration of the additional biological features associated with certain signatures. Such features should be quantified and formally accommodated in mathematical models, if methods for identification are to be further sharpened. At the same time, the pursuit of high-resolution mutational signatures by accounting for additional contextual features might be hindered by the limitations of current models. It can be argued that innovative models assuming niether complete mutual independence nor non-independence between the features of a signature could prove key to achieving the ideal compromise between flexibility and complexity that is warranted for powerful, stable and accurate delineation of mutational signatures.

As current and forthcoming approaches shed light on the mathematical properties of mutational signature discovery, the study of somatic mutation patterns will surely be extended through the addition of new signatures, aberration classes, contextual features, and previously unexamined cancer types. Meanwhile, the insights yielded by advances in this field will further our understanding of the causes, mechanisms and evolution of human malignancy, and provide new opportunities for cancer prevention and treatment.

## Key points

- The somatic mutations in a genome are the result of the activity of one or more mutational processes, some of which imprint a distinct mutational signature.
- Nonnegative matrix factorization (NMF) is the most widely used method for identifying mutational signatures.
- Alternative approaches include partly and fully probabilistic models, as well as NMF implementations offering greater ease of use.
- The study of mutational signatures can prove useful for cancer prevention and treatment efforts, including patient stratification and identification of novel mutagens.
- The field will likely be expanded with the inclusion of additional techniques, mutation classes, biological features and tumour types.

## Acknowledgments

We would like to thank Elizabeth Murchison and Ludmil Alexandrov for valuable discussions and critical advice.

## References

1. Stratton MR, Campbell PJ, Futreal PA. The cancer genome. Nature 2009; 458:719–724.

2. Alexandrov LB, Nik-Zainal S, Wedge DC, et al. Signatures of mutational processes in human cancer. Nature 2013; 500:415–421.

3. Vogelstein B, Papadopoulos N, Velculescu VE, et al. Cancer genome landscapes. Science 2013; 339:1546–1558.

4. Beerenwinkel N, Antal T, Dingli D, et al. Genetic progression and the waiting time to cancer. PLoS Comput. Biol. 2007; 3:e225.

5. Attolini CS-O, Michor F. Evolutionary theory of cancer. Ann. N. Y. Acad. Sci. 2009; 1168:23–51.

6. Yates LR, Campbell PJ. Evolution of the cancer genome. Nat. Rev. Genet. 2012; 13:795–806.

7. Alexandrov LB, Stratton MR. Mutational signatures: the patterns of somatic mutations hidden in cancer genomes. Curr. Opin. Genet. Dev. 2014; 24:52–60.

8. Roberts SA, Gordenin DA. Hypermutation in human cancer genomes: footprints and mechanisms. Nat. Rev. Cancer 2014; 14:786–800.

9. Pfeifer GP. Environmental exposures and mutational patterns of cancer genomes. Genome Med. 2010; 2:54.

10. Rubin AF, Green P. Mutation patterns in cancer genomes. Proc. Natl. Acad. Sci. U. S. A. 2009; 106:21766–21770.

11. Muller HJ. Further studies on the nature and causes of gene mutations. Proceedings of the 6th International Congress of Genetics 1932; 1:213–255.

12. Bauer H, Demerec M, Kaufmann BP. X-Ray Induced Chromosomal Alterations in Drosophila Melanogaster. Genetics 1938; 23:610–630.

13. Sax K. Chromosome Aberrations Induced by X-Rays. Genetics 1938; 23:494–516.

14. Howard BD, Tessman I. Identification of the altered bases in mutated single-stranded DNA: III. Mutagenesis by ultraviolet light. J. Mol. Biol. 1964; 9:372–375.

15. Pfeifer GP, You Y-H, Besaratinia A. Mutations induced by ultraviolet light. Mutat. Res. 2005; 571:19–31.

16. Setlow RB, Carrier WL. Pyrimidine dimers in ultraviolet-irradiated DNA’s. J. Mol. Biol. 1966; 17:237–254.

17. Govindan R, Ding L, Griffith M, et al. Genomic landscape of non-small cell lung cancer in smokers and never-smokers. Cell 2012; 150:1121–1134.

18. Pfeifer GP, Denissenko MF, Olivier M, et al. Tobacco smoke carcinogens, DNA damage and p53 mutations in smoking-associated cancers. Oncogene 2002; 21:7435–7451.

19. Harris RS. Cancer mutation signatures, DNA damage mechanisms, and potential clinical implications. Genome Med. 2013; 5:87.

20. Hunter C, Smith R, Cahill DP, et al. A hypermutation phenotype and somatic MSH6 mutations in recurrent human malignant gliomas after alkylator chemotherapy. Cancer Res. 2006; 66:3987–3991.

21. Helleday T, Eshtad S, Nik-Zainal S. Mechanisms underlying mutational signatures in human cancers. Nat. Rev. Genet. 2014; 15:585–598.

22. Alexandrov LB, Nik-Zainal S, Siu HC, et al. A mutational signature in gastric cancer suggests therapeutic strategies. Nat. Commun. 2015; 6:8683.

23. Fox EJ, Salk JJ, Loeb LA. Exploring the implications of distinct mutational signatures and mutation rates in aging and cancer. Genome Med. 2016; 8:30.

24. Li X, Wu WK, Xing R, et al. Distinct Subtypes of Gastric Cancer Defined by Molecular Characterization Include Novel Mutational Signatures with Prognostic Capability. Cancer Res. 2016; 76:1724–1732.

25. Poon SL, Huang MN, Choo Y, et al. Mutation signatures implicate aristolochic acid in bladder cancer development. Genome Med. 2015; 7:38.

26. Poon SL, McPherson JR, Tan P, et al. Mutation signatures of carcinogen exposure: genome-wide detection and new opportunities for cancer prevention. Genome Med. 2014; 6:24.

27. Secrier M, Li X, de Silva N, et al. Mutational signatures in esophageal adenocarcinoma define etiologically distinct subgroups with therapeutic relevance. Nat. Genet. 2016; 48:1131–1141.

28. Alexandrov LB, Jones PH, Wedge DC, et al. Clock-like mutational processes in human somatic cells. Nat. Genet. 2015; 47:1402–1407.

29. Nik-Zainal S, Davies H, Staaf J, et al. Landscape of somatic mutations in 560 breast cancer whole-genome sequences. Nature 2016; 534:47–54.

30. Schulze K, Imbeaud S, Letouzé E, et al. Exome sequencing of hepatocellular carcinomas identifies new mutational signatures and potential therapeutic targets. Nat. Genet. 2015; 47:505–511.

31. COSMIC: Signatures of Mutational Processes in Human Cancer. <u>http://cancer.sanger.ac.uk/cosmic/signatures</u> (27 April 2017, date last accessed).

32. Morganella S, Alexandrov LB, Glodzik D, et al. The topography of mutational processes in breast cancer genomes. Nat. Commun. 2016; 7:11383.

33. Nik-Zainal S, Alexandrov LB, Wedge DC, et al. Mutational processes molding the genomes of 21 breast cancers. Cell 2012; 149:979–993.

34. Alexandrov LB, Nik-Zainal S, Wedge DC, et al. Deciphering signatures of mutational processes operative in human cancer. Cell Rep. 2013; 3:246–259.

35. Shiraishi Y, Tremmel G, Miyano S, et al. A Simple Model-Based Approach to Inferring and Visualizing Cancer Mutation Signatures. PLoS Genet. 2015; 11:e1005657.

36. Lee DD, Seung HS. Learning the parts of objects by non-negative matrix factorization. Nature 1999; 401:788–791.

37. Paatero P, Tapper U. Positive matrix factorization: A non-negative factor model with optimal utilization of error estimates of data values. Environmetrics 1994; 5:111–126.

38. Brunet J-P, Tamayo P, Golub TR, et al. Metagenes and molecular pattern discovery using matrix factorization. Proc. Natl. Acad. Sci. U. S. A. 2004; 101:4164–4169.

39. Devarajan K. Nonnegative matrix factorization: an analytical and interpretive tool in computational biology. PLoS Comput. Biol. 2008; 4:e1000029.

40. Hutchins LN, Murphy SM, Singh P, et al. Position-dependent motif characterization using non-negative matrix factorization. Bioinformatics 2008; 24:2684–2690.

41. Pehkonen P, Wong G, Törönen P. Theme discovery from gene lists for identification and viewing of multiple functional groups. BMC Bioinformatics 2005; 6:162.

42. Xu M, Li W, James GM, et al. Automated multidimensional phenotypic profiling using large public microarray repositories. Proc. Natl. Acad. Sci. U. S. A. 2009; 106:12323–12328.

43. Nik-Zainal S, Van Loo P, Wedge DC, et al. The life history of 21 breast cancers. Cell 2012; 149:994–1007.

44. Lee DD, Seung HS. Algorithms for Non-negative Matrix Factorization. Advances in Neural Information Processing Systems 13 2001; 556–562.

45. Rousseeuw PJ. Silhouettes: A graphical aid to the interpretation and validation of cluster analysis. J. Comput. Appl. Math. 1987; 20:53–65.

46. Berry MW, Browne M, Langville AN, et al. Algorithms and applications for approximate nonnegative matrix factorization. Comput. Stat. Data Anal. 2007; 52:155–173.

47. Boutsidis C, Gallopoulos E. SVD based initialization: A head start for nonnegative matrix factorization. Pattern Recognit. 2008/4; 41:1350–1362.

48. Gehring JS, Fischer B, Lawrence M, et al. SomaticSignatures: inferring mutational signatures from single-nucleotide variants. Bioinformatics 2015; 31:3673–3675.

49. Gentleman RC, Carey VJ, Bates DM, et al. Bioconductor: open software development for computational biology and bioinformatics. Genome Biol. 2004; 5:R80.

50. Akre MK, Starrett GJ, Quist JS, et al. Mutation Processes in 293-Based Clones Overexpressing the DNA Cytosine Deaminase APOBEC3B. PLoS One 2016; 11:e0155391.

51. Durinck S, Stawiski EW, Pavía-Jiménez A, et al. Spectrum of diverse genomic alterations define non-clear cell renal carcinoma subtypes. Nat. Genet. 2015; 47:13–21.

52. Fei SS, Mitchell AD, Heskett MB, et al. Patient-specific factors influence somatic variation patterns in von Hippel-Lindau disease renal tumours. Nat. Commun. 2016; 7:11588.

53. Kovac M, Blattmann C, Ribi S, et al. Exome sequencing of osteosarcoma reveals mutation signatures reminiscent of BRCA deficiency. Nat. Commun. 2015; 6:8940.

54. Nagahashi M, Wakai T, Shimada Y, et al. Genomic landscape of colorectal cancer in Japan: clinical implications of comprehensive genomic sequencing for precision medicine. Genome Med. 2016; 8:136.

55. Ramakodi MP, Kulathinal RJ, Chung Y, et al. Ancestral-derived effects on the mutational landscape of laryngeal cancer. Genomics 2016; 107:76–82.

56. Weinhold N, Ashby C, Rasche L, et al. Clonal selection and double-hit events involving tumor suppressor genes underlie relapse in myeloma. Blood 2016; 128:1735–1744.

57. Ardin M, Cahais V, Castells X, et al. MutSpec: a Galaxy toolbox for streamlined analyses of somatic mutation spectra in human and mouse cancer genomes. BMC Bioinformatics 2016; 17:170.

58. Gaujoux R, Seoighe C. A flexible R package for nonnegative matrix factorization. BMC Bioinformatics 2010; 11:367.

59. Giardine B, Riemer C, Hardison RC, et al. Galaxy: a platform for interactive large-scale genome analysis. Genome Res. 2005; 15:1451–1455.

60. Goecks J, Nekrutenko A, Taylor J, et al. Galaxy: a comprehensive approach for supporting accessible, reproducible, and transparent computational research in the life sciences. Genome Biol. 2010; 11:R86.

61. Fischer A, Illingworth CJR, Campbell PJ, et al. EMu: probabilistic inference of mutational processes and their localization in the cancer genome. Genome Biol. 2013; 14:R39.

62. Banerjee A, Merugu S, Dhillon IS, et al. Clustering with Bregman Divergences. J. Mach. Learn. Res. 2005; 6:1705–1749.

63. Cemgil AT. Bayesian inference for nonnegative matrix factorisation models. Comput. Intell. Neurosci. 2009; 785152.

64. Févotte C, Cemgil AT. Nonnegative matrix factorizations as probabilistic inference in composite models. 2009 17th European Signal Processing Conference 2009; 1913–1917.

65. Schmidt MN, Winther O, Hansen LK. Bayesian Non-negative Matrix Factorization. Independent Component Analysis and Signal Separation 2009; 540–547.

66. Dempster AP, Laird NM, Rubin DB. Maximum Likelihood from Incomplete Data via the EM Algorithm. J. R. Stat. Soc. Series B Stat. Methodol. 1977; 39:1–38.

67. Weir BA, Woo MS, Getz G, et al. Characterizing the cancer genome in lung adenocarcinoma. Nature 2007; 450:893–898.

68. Schwarz G. Estimating the dimension of a model. Ann. Stat. 1978; 6:461–464.

69. Burnham KP, Anderson DR. Multimodel inference understanding AIC and BIC in model selection. Sociol. Methods Res. 2004; 33:261–304.

70. Kasar S, Kim J, Improgo R, et al. Whole-genome sequencing reveals activation-induced cytidine deaminase signatures during indolent chronic lymphocytic leukaemia evolution. Nat. Commun. 2015; 6:8866.

71. Kim J, Mouw KW, Polak P, et al. Somatic ERCC2 mutations are associated with a distinct genomic signature in urothelial tumors. Nat. Genet. 2016; 48:600–606.

72. Tan VYF, Févotte C. Automatic relevance determination in nonnegative matrix factorization with the β-divergence. IEEE Trans. Pattern Anal. Mach. Intell. 2013; 35:1592–1605.

73. Kingman JFC. Poisson Processes. Encyclopedia of Biostatistics 2005.

74. Rosales RA, Drummond RD, Valieris R, et al. signeR: an empirical Bayesian approach to mutational signature discovery. Bioinformatics 2017; 33:8–16.

75. Casella G. Empirical Bayes Gibbs sampling. Biostatistics 2001; 2:485–500.

76. Altman NS. An Introduction to Kernel and Nearest-Neighbor Nonparametric Regression. Am. Stat. 1992; 46:175–185.

77. Krawczak M, Ball EV, Cooper DN. Neighboring-nucleotide effects on the rates of germ-line single-base-pair substitution in human genes. Am. J. Hum. Genet. 1998; 63:474–488.

78. Pleasance ED, Cheetham RK, Stephens PJ, et al. A comprehensive catalogue of somatic mutations from a human cancer genome. Nature 2010; 463:191–196.

79. Pritchard JK, Stephens M, Donnelly P. Inference of population structure using multilocus genotype data. Genetics 2000; 155:945–959.

80. Blei DM, Ng AY, Jordan MI. Latent Dirichlet Allocation. J. Mach. Learn. Res. 2003; 3:993–1022.

81. Ding C, Li T, Peng W. On the equivalence between Non-negative Matrix Factorization and Probabilistic Latent Semantic Indexing. Comput. Stat. Data Anal. 2008; 52:3913–3927.

82. Blokzijl F, de Ligt J, Jager M, et al. Tissue-specific mutation accumulation in human adult stem cells during life. Nature 2016; 538:260–264.

83. Zámborszky J, Szikriszt B, Gervai JZ, et al. Loss of BRCA1 or BRCA2 markedly increases the rate of base substitution mutagenesis and has distinct effects on genomic deletions. Oncogene 2017; 36:746–755.

84. Schneider TD, Stephens RM. Sequence logos: a new way to display consensus sequences. Nucleic Acids Res. 1990; 18:6097–6100.

85. Cleveland WS, McGill R. Graphical Perception: Theory, Experimentation, and Application to the Development of Graphical Methods. J. Am. Stat. Assoc. 1984; 79:531–554.

86. Rosenthal R, McGranahan N, Herrero J, et al. DeconstructSigs: delineating mutational processes in single tumors distinguishes DNA repair deficiencies and patterns of carcinoma evolution. Genome Biol. 2016; 17:31.

87. Bruna A, Rueda OM, Greenwood W, et al. A Biobank of Breast Cancer Explants with Preserved Intra-tumor Heterogeneity to Screen Anticancer Compounds. Cell 2016; 167:260–274.e22.

88. Goh G, Schmid R, Guiver K, et al. Clonal Evolutionary Analysis during HER2 Blockade in HER2-Positive Inflammatory Breast Cancer: A Phase II Open-Label Clinical Trial of Afatinib+/-Vinorelbine. PLoS Med. 2016; 13:e1002136.

89. Hao J-J, Lin D-C, Dinh HQ, et al. Spatial intratumoral heterogeneity and temporal clonal evolution in esophageal squamous cell carcinoma. Nat. Genet. 2016; 48:1500–1507.

90. Kanu N, Cerone MA, Goh G, et al. DNA replication stress mediates APOBEC3 family mutagenesis in breast cancer. Genome Biol. 2016; 17:185.

91. Murchison EP, Wedge DC, Alexandrov LB, et al. Transmissible dog cancer genome reveals the origin and history of an ancient cell lineage. Science 2014; 343:437–440.

92. Rahbari R, Wuster A, Lindsay SJ, et al. Timing, rates and spectra of human germline mutation. Nat. Genet. 2016; 48:126–133.

93. Davies H, Glodzik D, Morganella S, et al. HRDetect is a predictor of BRCA1 and BRCA2 deficiency based on mutational signatures. Nat. Med. 2017.

94. Alexandrov LB, Ju YS, Haase K, et al. Mutational signatures associated with tobacco smoking in human cancer. Science 2016; 354:618–622.

95. Hollstein M, Alexandrov LB, Wild CP, et al. Base changes in tumour DNA have the power to reveal the causes and evolution of cancer. Oncogene 2017; 36:158–167.

96. Loeb LA, Springgate CF, Battula N. Errors in DNA replication as a basis of malignant changes. Cancer Res. 1974; 34:2311–2321.

97. Behjati S, Gundem G, Wedge DC, et al. Mutational signatures of ionizing radiation in second malignancies. Nat. Commun. 2016; 7:12605.

98. Gao Y, Church G. Improving molecular cancer class discovery through sparse non-negative matrix factorization. Bioinformatics 2005; 21:3970–3975.

99. Guan N, Huang X, Lan L, et al. Graph Based Semi-supervised Non-negative Matrix Factorization for Document Clustering. 2012 11th International Conference on Machine Learning and Applications 2012; 1:404–408.

100. Hillebrand M, Kreßel U, Wöhler C, et al. Traffic Sign Classifier Adaption by Semi-supervised Co-training. Artificial Neural Networks in Pattern Recognition 2012; 193–200.

101. Lefevre A, Bach F, Févotte C. Semi-supervised NMF with time-frequency annotations for single-channel source separation. ISMIR 2012: 13th International Society for Music Information Retrieval Conference 2012.

102. Morikawa Y, Yukawa M. A sparse optimization approach to supervised NMF based on convex analytic method. 2013 IEEE International Conference on Acoustics, Speech and Signal Processing 2013; 6078–6082.

103. Peharz R, Pernkopf F. Sparse nonnegative matrix factorization with ℓ0-constraints. Neurocomputing 2012; 80:38–46.

104. Sindhwani V, Ghoting A. Large-scale Distributed Non-negative Sparse Coding and Sparse Dictionary Learning. Proceedings of the 18th ACM SIGKDD International Conference on Knowledge Discovery and Data Mining 2012; 489–497.

105. Zheng C-H, Huang D-S, Sun Z-L, et al. Nonnegative independent component analysis based on minimizing mutual information technique. Neurocomputing 2006/3; 69:878–883.

106. Chen M, Chen W-S, Chen B, et al. Non-negative Sparse Representation Based on Block NMF for Face Recognition. Biometric Recognition 2013; 26–33.

